# Decoupled phenology and PSII thermal plasticity in seasonally dry tropical forest trees

**DOI:** 10.64898/2026.06.10.731154

**Authors:** Rakesh Tiwari, Pratibha T. Bhagawad, H Shiva Naik, Rakshit G. Hosamani, N Prakash, M. S. Jayanath Babu, B.S. Shreeloka, M. M. Madhur, G Sagara, Tharun Naik, Amani Khanum Soor, N Vijayashree, B.S Lavanya, T. G Kishore, N Madhusudhana, Mahesh K Patil, Nanda Appaji, K.G. Somashekhar Achar, SH Basavarajappa, Robert Muscarella, Yogendra Kambalagere

## Abstract

We tested whether seasonal plasticity in photosystem II (PSII) heat tolerance aligns with leaf phenology in tropical trees, comparing evergreen and deciduous species across the wet⍰to⍰post⍰wet transition in a seasonally dry tropical forest of the central Western Ghats, India. This transition, preceding drought⍰induced senescence, represents the key window to assess true unstressed thermal plasticity. PSII thermal traits: damage onset (*T*_5_), damage midpoint (*T*_50_), and temperature between damage onset to full loss (*T*_95_–*T*_5_), decline width (DW) were quantified in 27 co⍰occurring species during the wet (27.5□°C) and post⍰wet (31.6□°C) periods. Contrary to phenology⍰based predictions, PSII plasticity was not structured by leaf habit or successional status. Both *T*_5_ (+1.7□°C) and *T*_50_ (+0.9□°C) increased significantly across seasons, but responses were species⍰specific, with evergreen and deciduous trees acclimating similarly. The prevention⍰versus⍰forbearance trade⍰off (*T*_5_ – DW relationship) remained conserved, though leaf habits diverged under post⍰wet conditions. Thermal safety margins based on *T*_50_ were large, but *T*_5_ revealed vulnerable late⍰successional evergreens (*Saraca asoca, Ficus* spp.) and *Careya arborea*. These results show that PSII thermotolerance regulation operates largely independently of drought⍰avoidance phenology, indicating species identity and not leaf habit drives plasticity in PSII thermal response in seasonally dry tropical forests.

## Introduction

Tropical and subtropical forests globally are approaching their operational thermal limits, with climate warming projected to push many species beyond their historical temperature thresholds within this century (Doughty *et al*. 2023). Photosystem II (PSII) thermal damage characterized by the loss of photosynthetic quantum yield represents a critical ecophysiological measure of tree performance, particularly in seasonally variable environments where temperature, water availability and vapor pressure deficit (VPD) vary starkly across the annual cycle (Kattge *et al*. 2011; Sastry & Barua 2017; Dusenge, Duarte & Way 2019). Understanding when and how tree leaves experience thermal stress, and the physiological capacity to acclimate or tolerate this stress, is hence essential for predicting species responses to warming.

Despite growing evidence that PSII thermal tolerance is linked to habitat and life⍰history strategy—such that lowland species and those with higher leaf mass per area tend to exhibit higher T50 values (Sastry & Barua 2017; Slot *et al*. 2021), and thermotolerance differs between evergreen and deciduous species (Sastry & Barua, 2017) - most tropical studies remain constrained to single seasons or phenological extremes. This limits understanding of how seasonal plasticity in PSII stability unfolds across the productive growing season, particularly in the post⍰wet period when temperatures are rising but drought and peak heat stress have not yet peaked (Tiwari *et al*. 2025).

Most of the studies of PSII thermal tolerance in tropics have been conducted either during the wet period, when ambient temperatures are lower, soils are saturated, and VPD is moderate, but leaves are mature or young and experience low stress (Dusenge *et al*. 2019; Slot *et al*. 2021); or during the peak dry season, when temperatures and VPD are at their extremes, but many deciduous species are already partially or fully defoliated, or leaves are senescing and under severe water stress, while in some species, new leaf flushes emerge post-summer (Nanda, Prakasha, Murthy & Suresh 2012; Sastry & Barua 2017; Sastry, Guha & Barua 2018; Silva, Espírito-Santo, Santos & Rodrigues 2020; Ramos *et al*. 2023). Thus, the focus on two extreme periods (Sastry & Barua 2017; Tiwari *et al*. 2021), has led to the gap in understanding of the physiologically critical intermediate state: the post-wet (pre-senescence) period - the period when leaves are fully mature and capable of photosynthetic function, water availability is still favourable, and temperatures and VPD are rising but not yet extreme. This window represents the true peak canopy functional period for deciduous trees in the seasonally dry tropical forests (SDTF) when they are most productive but also when thermal stress and dehydration risk are first beginning to accumulate (Guan *et al*. 2015; Tiwari *et al*. 2025). Yet, PSII heat tolerance of fully mature leaves during this post-wet period remains largely unquantified in seasonally dry tropical forests.

Leaf longevity integrates structural robustness and phenological strategy; long-lived leaves evolve enhanced membrane stability and repair under chronic exposure, while short-lived ones favour high initial productivity over extreme tolerance (Russo & Kitajima 2016; Slot *et al*. 2021). Deciduous phenology not only avoids peak heat but modulates PSII traits during peak productivity windows, contrasting evergreen constitutive tolerance (Sastry *et al*. 2018). Especially in seasonally dry tropical forests, co-occurring tree species exhibit diverse phenological strategies, from drought-deciduous species that inherently avoid peak-summer heat to others that tolerate warmer conditions - setting the stage for contrasting investments in PSII thermal tolerance during non-extreme periods (Tiwari *et al*. 2025).

A widely held but rarely tested assumption follows from this phenological contrast: that leaf habit should predict the degree of seasonal PSII thermal plasticity. Evergreen species, retaining leaves year-round and experiencing the full thermal amplitude of the annual cycle, are expected to regulate PSII thermotolerance continuously upregulating membrane stability, D1 turnover, and heat-shock protein synthesis as temperatures rise, and modulating these investments seasonally in response to growth temperature (Yamori, Hikosaka & Way 2014; Slot *et al*. 2021). Deciduous species, by contrast, are expected to invest less in physiological thermotolerance regulation precisely because phenological avoidance pre-empts the need: if leaves are shed before peak heat stress arrives, the energetic cost of maintaining PSII stability under extreme temperatures is avoided entirely (Sastry & Barua 2017; Sastry *et al*. 2018). Under this framework, seasonal plasticity in PSII tolerance should be greater in evergreens than deciduous species, and leaf habit should therefore serve as a reliable functional proxy for a species’ capacity to physiologically buffer warming. This assumption has informed trait-based approaches to predicting tropical forest responses to climate warming (Doughty *et al*. 2023), yet it has not been directly tested across a diverse community of co-occurring species measured under matched conditions during a non-stress seasonal transition, precisely the context needed to isolate thermally-driven plasticity from the confounding effects of drought stress and leaf senescence that characterise peak dry-season comparisons.

PSII heat stability, quantified by the temperatures at which *F*_v_/*F*_m_ declines by 5% (*T*_5_, onset of stress) and 50% (*T*_50_, irreversible damage), reflects the integrity of reaction-centre proteins, thylakoid membranes, and supporting mechanisms including lipid composition, chloroplast small heat-shock proteins, and the D1 repair cycle (Schreiber & Berry 1977; Schreiber 2007; O’sullivan *et al*. 2017; Tiwari *et al*. 2021; Slot *et al*. 2021). Maintaining PSII under heat imposes energetic costs via protein turnover and repair, which acclimate to growth temperature but remain biochemically constrained, yielding modest *T*_50_ plasticity that rarely fully compensates seasonal thermal shifts (Yi *et al*. 2022). The decline width (*DW, T*_95_ – *T*_5_) captures response dynamics beyond absolute thresholds: narrow DW signals abrupt PSII failure despite high *T*_5_ (prevention-focused), while wide DW indicates buffered decline post-onset (repair/forbearance-focused), revealing distinct thermal strategies among species (Tiwari *et al*. 2021).

PSII thermal tolerance is expected to be partially plastic because the integrity of the reaction centre, thylakoid membranes, and associated repair machinery are energetically and biochemically modifiable traits. Elevated growth temperature can enhance PSII heat tolerance by upregulating the synthesis and turnover of the D1 protein, stimulating chloroplast small heat⍰shock proteins, and adjusting lipid composition and non⍰photochemical quenching components, all of which stabilise PSII under heat (Yamori *et al*. 2014; Tiwari *et al*. 2021; Posch *et al*. 2026). However, each of these adjustments imposes costs in nitrogen, protein synthesis, and ATP that compete with investment in photosynthetic capacity and growth, implying that species will strategically balance constitutive thermal robustness against acclimation potential based on habitat thermal variability and leaf economics (Sastry & Barua 2017; Tiwari *et al*. 2021, 2026).

Given the typically modest PSII thermal acclimation observed in tropical systems (Tiwari *et al*. 2021; Posch *et al*. 2026), a ~3–4□°C seasonal warming from wet to post⍰wet periods in the Western Ghats (Krishnadas, Kumar & Comita 2016; Tiwari *et al*. 2025) implies only partial compensation via *T*_5_ and *T*_50_ shifts, but the extent of this seasonal plasticity, particularly for fully mature leaves of diverse evergreen and deciduous species, has not yet been quantified. Empirical field and urban garden studies across diverse biomes show that PSII heat⍰tolerance metrics (*T*_5_, *T*_50_) generally acclimate modestly to growth temperature and to seasonal shifts, but this plasticity is partial, species⍰ and strategy⍰dependent, and rarely fully compensates for thermal regime shifts under climate warming (Sastry & Barua 2017; Slot *et al*. 2021; Tarvainen *et al*. 2022; Kullberg & Feeley 2024; Manzi *et al*. 2025). These studies also highlight that leaf habit, economic traits, and drought exposure modulate the degree of acclimation, yet most remain constrained spatially and temporally, with few datasets tracking the same individuals across phenological stages and under the same canopy microclimates.

Seasonally dry forests of the central Western Ghats provide an ideal setting to address these questions, combining pronounced monsoon–post-monsoon transitions (~4°C warming from wet to post-wet) with co-occurring evergreen, semi-evergreen, and deciduous species spanning the slow–fast spectrum (Krishnadas *et al*. 2016; Tiwari *et al*. 2025). Here we quantify PSII heat tolerance (*T*_5_, *T*_50_, *DW*), and leaf habit for 27 species across wet and post-wet periods, outside peak hot–dry stress to test seasonal plasticity during physiologically critical windows. We answer the following questions:

1. Is PSII thermal tolerance (*T*_5_, *T*_50_, *DW*) plastic between wet and post-wet periods, and does this differ between evergreen/deciduous habits or along the successional spectrum?
2. Do seasonal/habit differences reveal distinct thermal safety strategies relevant to warming vulnerability in seasonally dry tropical forests?

Building on the assumption that phenological strategy predicts PSII thermal plasticity, we predict that deciduous species will show limited seasonal upregulation of *T*_5_ and *T*_50_ across the wet-to-post-wet transition, consistent with a phenological avoidance strategy that renders physiological thermotolerance regulation unnecessary during this window; while evergreen species, lacking the option of leaf shedding, will show greater seasonal plasticity as a compensatory response to rising temperatures (Sastry & Barua 2017; Slot *et al*. 2021; Kullberg & Feeley 2024). We further predict that the *T*_5_ ∝ *DW* prevention-forbearance trade-off will be conserved across seasons, reflecting a stable axis of PSII thermal strategy that is independent of seasonal acclimation, and that thermal safety margins will be narrower in evergreens under post-wet warming given their year-round leaf retention and continuous exposure to rising temperatures. Testing these predictions across 27 co-occurring species in a non-stress seasonal window where thermal signal is not confounded by drought or senescence provides a uniquely clean context for evaluating whether phenological strategy and PSII plasticity are coupled or independent dimensions of thermal adaptation in seasonally dry tropical forests.

## Materials and methods

### Study site

The study was conducted in a seasonally dry tropical forest within Bhadra Reserve Forest, Shivamogga district, Karnataka, India (13.747462° N, 75.611980° E; 680 m asl). The site features predominantly deciduous tree species interspersed with some evergreens (Krishnamurthy *et al*. 2010). It experiences a 5–6-month monsoon (~1200 mm rainfall), followed by a 3–4-month winter with rising daytime temperatures and declining nighttime temperatures (Figure 1). Notably, maximum air temperatures during the wet period (June–August) average 4°C lower than in the post-wet period (October–December), when daytime highs increase while nighttime lows decrease. The wet period (June–August) had a mean maximum air temperature of 27.5□±□0.2□°C. The post-wet period (October–December) was 4.1□°C warmer, with a mean maximum temperature of 31.6□±□0.14□°C. The summer period recorded the highest mean maximum temperature of 35.8□±□0.17□°C, 4.2□°C higher than the post-wet period (Figure 1).

**Figure 1.**
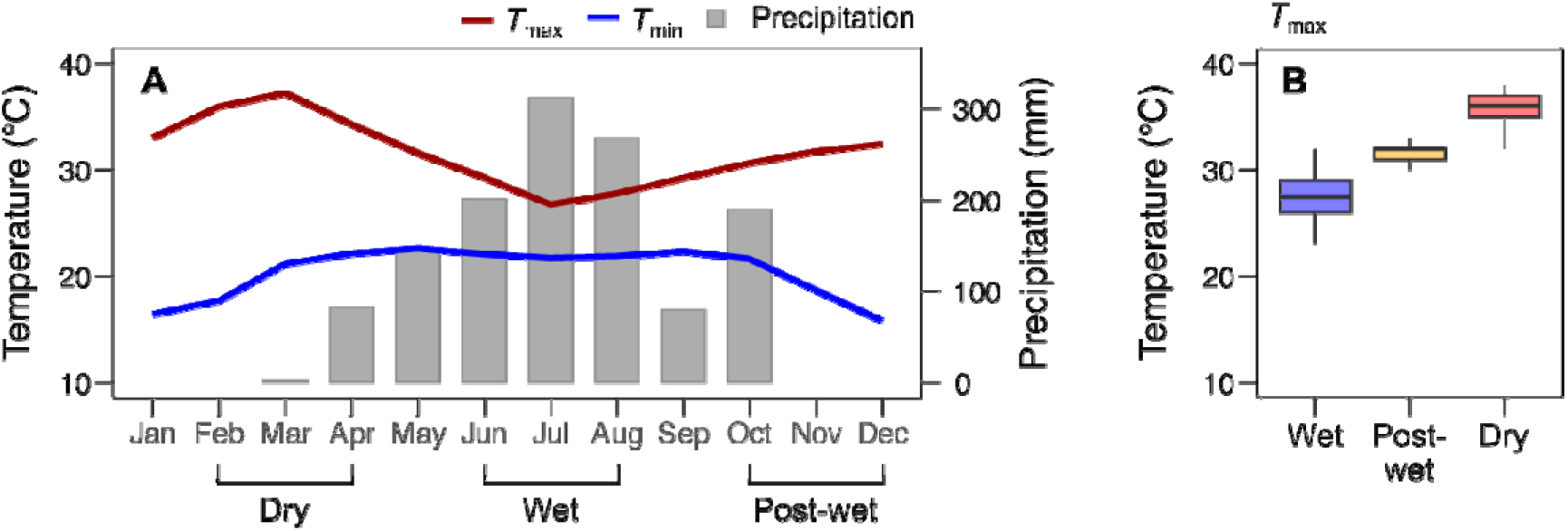
Panel (A) Seasonal monthly mean air temperature maxima/minima, and monthly total precipitation at Singanamane weather station (13.719217°N, 75.634778°E), located 2 km from the study site. Data cover March 2025–February 2026 (annual precipitation: 1284 mm). Panel B shows maximum air temperature ranges during the three periods.

### Plant materials

We selected the most dominant canopy and subcanopy tree species spanning a spectrum of leaf longevity—from short7llived leaves that persisted only through the wet and post7lwet periods to evergreens with year⍰round foliage, based on long-term phenological data from Nanda (2012) for the same site. This set included drought⍰deciduous heat⍰avoiders as well as typical evergreens with longer leaf longevity, thereby encompassing diverse leaf phenological and thermal adaptation strategies.

The study includes a total of 27 tree species characteristic of seasonally dry tropical forests in the central Western Ghats, Karnataka, India, spanning varied successional stages (Supplementary Table S1) (Pascal 1988). Species composition shifted from predominantly deciduous in drier stands to semi⍰evergreen and evergreen in wetter forest types along rainfall and topographic gradients. The site we studied is dominated by deciduous species such as *Xylia xylocarpa, Terminalia paniculata, Lannea coromandelica*, and *Careya arborea*, with *Terminalia paniculata* and *Xylia xylocarpa* jointly accounting for >50% of basal area (Krishnamurthy *et al*. 2010), alongside *Tectona grandis, Catunaregam spinosa*, and *Terminalia anogeissiana*. Typical evergreen species include *Artocarpus heterophyllus, Schleichera oleosa, Cassine glauca, Ficus benghalensis, F. macrocarpa, F. religiosa*, and *Santalum album*, among others. The canopy is about 15m tall and has about two strata. Detailed taxonomy, families, and ecological attributes including succession stage and canopy properties are listed in Supplementary Table S1. For each species, we sampled five individual trees (biological replicates). The site is further characterized by an intense herbivore and carnivore fauna that strongly suppresses understorey growth.

We conducted two measurement campaigns: (a) the late summer monsoon period (mid-August to late September 2025), referred to as the ‘wet period’, and (b) the second-to-fourth week of November 2025, post-rainfall and characterized by low minimum air temperatures and rising maximum air temperatures, referred to as the ‘post-wet period’. For both campaigns, we sampled leaves representing the predominant phenological stage at each time point – we note that in all species bore mature leaves during both the campaigns. Leaf samples were collected from about 4-5m from above ground using telescopic pruners. In both campaigns, leaves that were exposed to full sunlight were collected before 9AM. Samples were transported to lab in dark polythene bags under hydrated conditions.

### Heat treatment assay

We adopted Tiwari et al., (2021) and Krause (2010) protocols. We cut leaf 20 mm diameter leaf discs underwater, choosing the major lamina part of the leaf – avoiding midrib. For small leaves, we excised one disc per leaves and for the large lamina leaves, we excised 1-3 discs per leaf. Each assay consisted of at least 10 individual leaf discs from a tree. Leaf discs were randomly selected and wrapped in a layer of moist tissue paper on both sides covering the cut edge of the leaf discs. Care was taken not to touch the leaves, and the leaf discs always remained in water or covered in damp tissue paper to avoid anaerobiosis (Harris & Heber 1993). The discs were transferred into separate plastic bags (50 × 60 mm) keeping them flat and covered in a thin film of water. The bags were subjected to heat treatment in sets of at least three discs per temperature point. Heat treatment was conducted in temperature-controlled thermos flasks with pre-heated water. The water temperature was recorded in the central area of the flask using a calibrated K-type thermocouple probe. Separate sets of leaf discs were treated to one of the following temperatures points: 35, 40, 42, 44, 46, 48, 50, 52, and 60°C and a set of untreated discs were used as controls (about 25 °C). After heat treatment, the leaf discs were transferred to petri-dishes with tissue paper and water. The Petri dishes were incubated in the lab condition for 24h. After incubation period, the discs were tap-dried, wrapped in aluminium envelopes for dark adaptation followed by *F*_v_/*F*_m_ measurements using the Multispeq v2.0 (PhotosynQ, Michigan State University, USA)(Kuhlgert *et al*. 2016).

### Determination of PSII thermal tolerance parameters

A three-parameter logistic curve was fitted to each *F*_v_/*F*_m_ - temperature response series using the following equation:

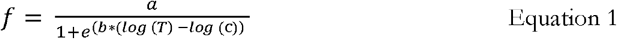

In this equation, *T* is defined as the treatment temperature, *b* is the steepness of the curve, *a* is the upper asymptote, and *c* is the inflection point. From the fits, we extracted (i) *T*_5_, the temperature at which *F*_v_/*F*_m_ declines to 5% of the maximum level and (ii) *T*_50_, the temperature where *F*_v_/*F*_m_ is half of the maximum. In addition, *T*_95_ was determined. This parameter is defined as the temperature at which the values of *F*_v_/*F*_m_ decrease to 95% of the maximum level. The differences between *T*_5_ and *T*_95_ are henceforward referred to as the decline width (*DW* = *T*_95_–*T*_5_), which is defined as the temperature range in which *F*_v_/*F*_m_ declined from 95% to 5% of the maximum level (Tiwari *et al*. 2021).

### Data analysis

PSII thermal tolerance parameters (*T*_5_, *T*_50,_ *DW* = *T*_95_ – *T*_5_) were derived from three-parameter logistic fits to Fv/Fm-temperature res ponse curves (Equation 1). Species-level plasticity (Δ*T*_5_ = post-wet – wet *T*_5_; similarly, Δ*T*_50_) was modelled using linear mixed-effects models (lmer; Bates *et al*. (2015)) with leaf habit (evergreen/deciduous), season (wet/post-wet), and their interaction as fixed effects, species as random intercept, (lme4 package). Type III ANOVA with Satterthwaite’s degrees of freedom tested fixed effects (lmerTest), and marginal *R*^2^ quantified habit effects (MuMIn package Bartoń (2018)) (Supplementary Table S2). Overall community means used analogous intercept-only models. Relationships among Δ*T*_5_–Δ*T*_50_ and Δ*T*_5_–DW employed SMA regression (smatr package) and concordance tests. Thermal safety margins (TSM_50_ = Δ*T*_50_ – *T*_air (90th)_; similarly, *TSM*_5_ = Δ*T*_5_ – *T*_air (90th)_) were compared via mixed models. Analyses used R 4.5.2 1 (R Core Team, 2025); figures via ggplot2 (Wickham 2009).

## Results

### Wet to post-wet period plasticity in PSII thermal tolerance

PSII thermal tolerance onset (*T*_5_) increased significantly from wet to post-wet periods (*F*_(1, 241)_ = 9.24, *p* = 0.0026), indicating seasonal plasticity, though this simple model explained only 2.7% of variance (marginal *R*^2^) (Figure 2, A & C). A species × season interaction model revealed strong interspecific variation (*F*_(27, 215)_ = 3.69, *p* < 0.0001) and confirmed the seasonal increase (*F*_(1, 215)_ = 10.14, *p* = 0.0017), with a significant species × season interaction (*F*_(26, 215)_ = 1.71, *p* = 0.021), indicating that the magnitude of *T*_5_ plasticity varied substantially among species (marginal *R*^2^ = 0.36). These results demonstrate that while *T*_5_ exhibits clear seasonal acclimation, species differ markedly in their plastic response to warming. Neither leaf habit nor successional status significantly modulated *T*_5_ plasticity across seasons (habit : season *F*_(1, 240)_ = 0.0025, p = 0.96; succession : season *F*_(2, 239)_ = 0.13, *p* = 0.88), with the main seasonal effect persisting (*F*_(1, 240)_ = 9.18, *p* = 0.0027) but explaining limited variance (*R*^2^ < 7%) (Supplementary Figure S1A). Wet-to-post-wet changes in *T*_5_, Δ*T*_5_ did not differ between deciduous and evergreen species (*F*_(1, 240)_ = 0.009, *p* = 0.93), and leaf habit explained essentially none of the variance in plasticity (Figure 3, A).

**Figure 2.**
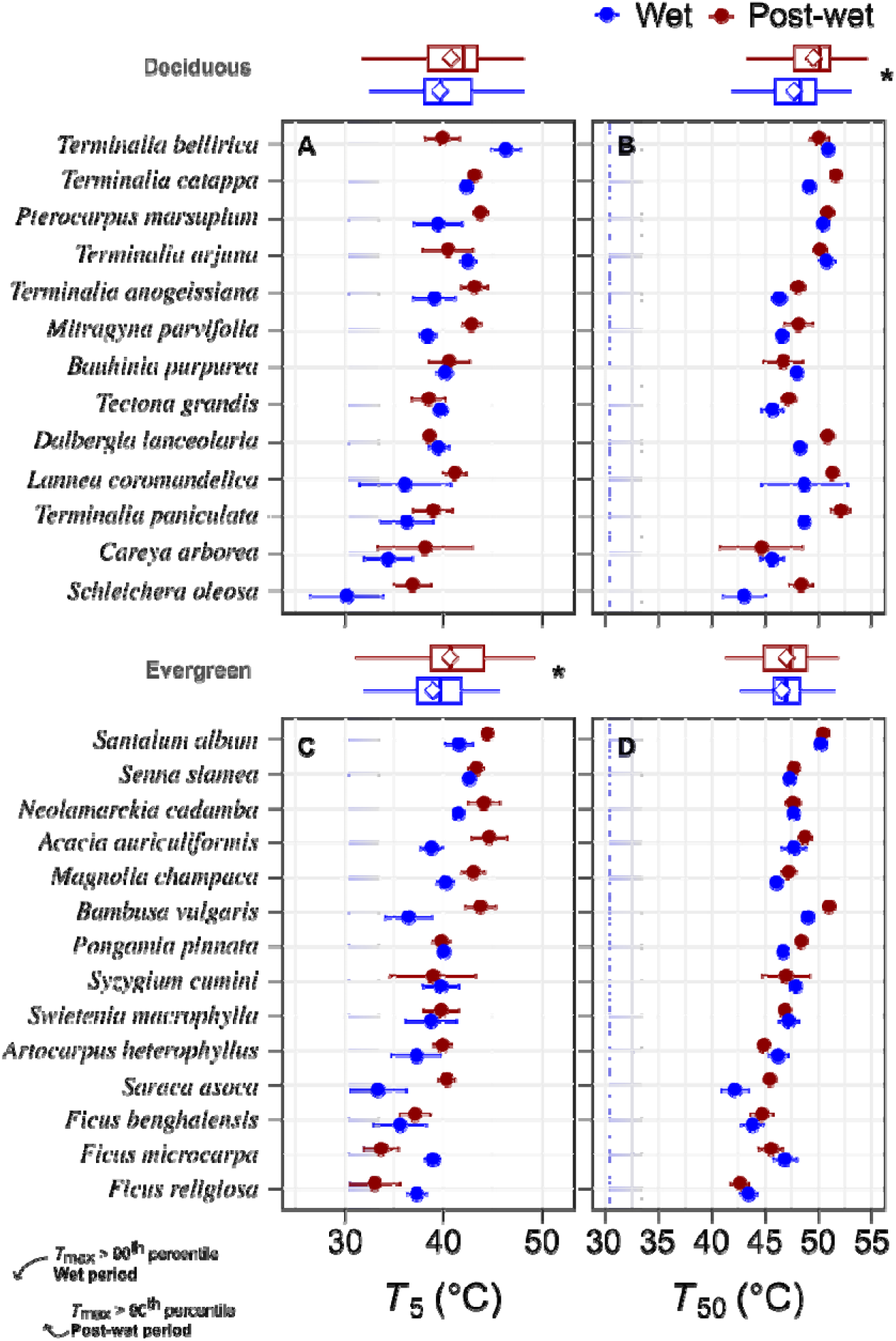
PSII thermal tolerance (*T*_5_ and *T*_50_) of co-occurring deciduous and evergreen tree species during wet monsoon and post-wet periods, relative to seasonal air temperature maxima (*T*_max_ > 90^th^ percentile). Boxplots depict *T*_5_ (left panels, A,C) and *T*_50_ (right panels, B,D) thresholds for leaf heat tolerance in deciduous (top row) and evergreen (middle row) species. *T*_5_ marks 5% *F*_v_/*F*_m_ decline (functional onset), *T*_50_ marks 50% decline (damage midpoint). Bottom panels show *T*_max_ distributions (30.4°C wet; 33.4°C post-wet), revealing most species retain large *T*_50_ buffers while select evergreens (*Saraca asoca, Ficus* spp.) and Careya arborea operate near *T*_5_ functional limits.

**Figure 3.**
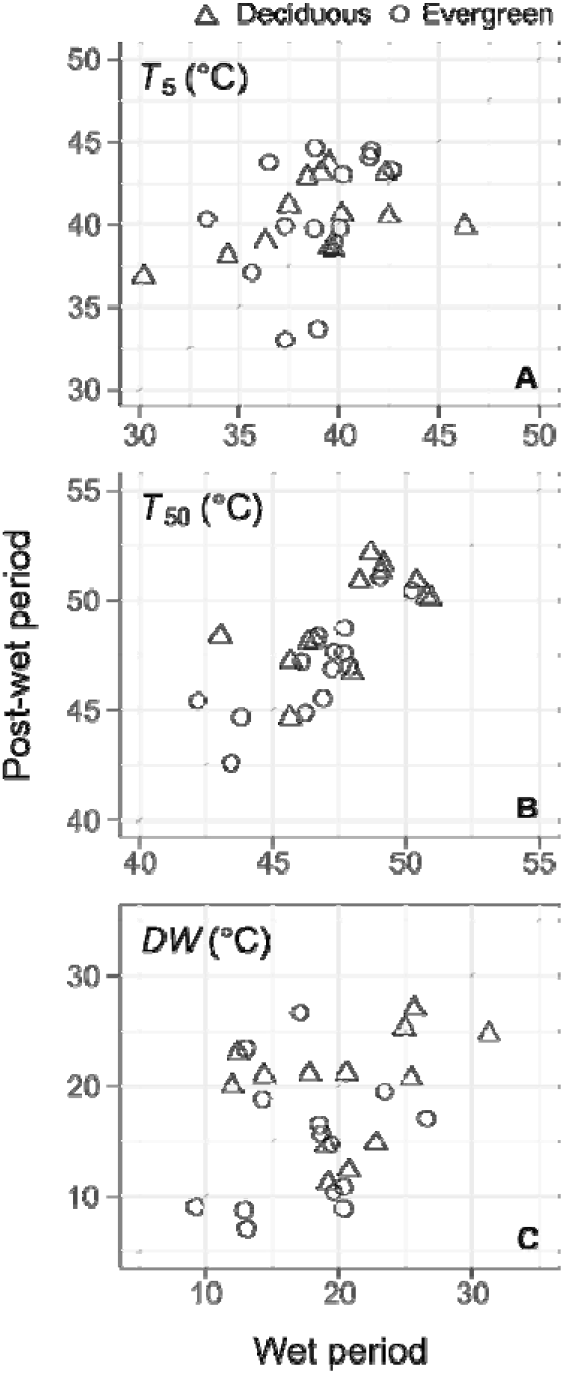
Wet-to-post-wet period plasticity in PSII thermal tolerance parameters (*T*_5_, *T*_50_ and decline width, panels A-C) across 27 tree species in a seasonally dry tropical forest, Central Western Ghats, India. Circles represent evergreen species; triangles represent deciduous species. Each symbol shows species mean (n=5 trees). Dashed line indicates 1:1 relationship.

PSII thermal damage midpoint (*T*_50_) increased significantly from wet to post-wet periods (*F*_(1, 241)_ = 8.61, *p* = 0.0037), indicating seasonal plasticity, though this simple model explained only 1.8% of variance (marginal *R*^2^) (Figure 2, B & D). A species × season interaction model revealed strong interspecific variation (*F*_(26, 122)_ = 8.19, *p* < 0.0001) and confirmed the seasonal increase (*F*_(1, 122)_ = 9.16, *p* = 0.0030), with a non-significant species × season interaction (*F*_(26, 115)_ = 1.28, *p* = 0.19), indicating limited species-level variation in the magnitude of *T*_50_ plasticity (marginal *R*^2^ = 0.49). Leaf habit showed a main effect (*F*_(1, 25)_ = 4.76, *p* = 0.039) but no interaction with season (habit : season *F*_(1, 240)_ = 2.60, *p* = 0.11), while successional status marginally modulated seasonal plasticity (succession : season *F*_(2, 239)_ = 2.79, *p* = 0.064), with the main seasonal effect persisting (*F*_(1, 239)_ = 5.04, *p* = 0.026) but explaining limited variance (*R*^2^ < 10%) (Supplementary Figure S1B). These results demonstrate that while *T*_50_ exhibits modest seasonal acclimation, plasticity magnitude is largely consistent across species, with weak modulation by leaf habit and a marginal influence of successional status. Wet-to-post-wet changes in *T*_50_, Δ*T*_50_ did not differ between deciduous and evergreen species (*F*_(26, 115)_ = 2.14, *p* = 0.16), with leaf habit (Figure 3, B).

PSII decline width (*DW*) showed a marginal increase from wet to post-wet periods (*F*_(1, 241)_ = 3.10, *p* = 0.079), suggesting weak seasonal plasticity, with this simple model explaining only 1.0% of variance (marginal *R*^2^). A species × season interaction model revealed strong interspecific variation (*F*_(26, 215)_ = 3.11, *p* < 0.0001) and confirmed the marginal seasonal trend (*F*_(1, 215)_ = 3.39, *p* = 0.067), with a significant species × season interaction (*F*_(26, 215)_ = 1.64, p = 0.031), indicating species differed substantially in DW plasticity magnitude (marginal *R*^2^ = 0.32). Leaf habit showed a main effect (F_(1, 25)_ = 5.99, *p* = 0.022) but no interaction with season (habit : season *F*_(1, 240)_ = 0.91, *p* = 0.34), while successional status showed no modulation of seasonal plasticity (succession : season *F*_(2, 239)_ = 1.49, *p* = 0.23), with the main seasonal effect persisting marginally (*F*_(1, 239)_ = 4.66, *p* = 0.032) but explaining limited variance (*R*^2^ < 6%). These results demonstrate that while *DW* exhibits weak seasonal plasticity, species differ markedly in their response, though neither habit nor successional status significantly modulates this plasticity. Wet-to-post-wet changes in *DW*, Δ*DW* did not differ between deciduous and evergreen species (*F*_(26, 115)_ = 0.65, *p* = 0.43), with leaf habit explaining only 0.7% of the variance in plasticity (Figure 3, C).

### Coordinated seasonal shifts in *T*_5_ and *T*_50_

Post-wet and wet period species mean seasonal changes in *T*_5_ and *T*_50_ were moderately correlated (r = 0.59, 95% CI: 0.24–0.80, p = 0.0023). A sign⍰based test of concordance indicated that most species exhibited parallel shifts in *T*_5_ and *T*_50_. Fisher’s exact test suggested that species with positive Δ*T*_5_ (with higher valued during post-wet period) were substantially more likely to also show positive Δ*T*_50_ (odds ratio = 7.3, 95% CI: 0.82–106), although this association was only marginally significant (p = 0.061). These results indicate that *T*_5_ and *T*_50_ tend to change in similar directions across seasons (Figure 4).

**Figure 4.**
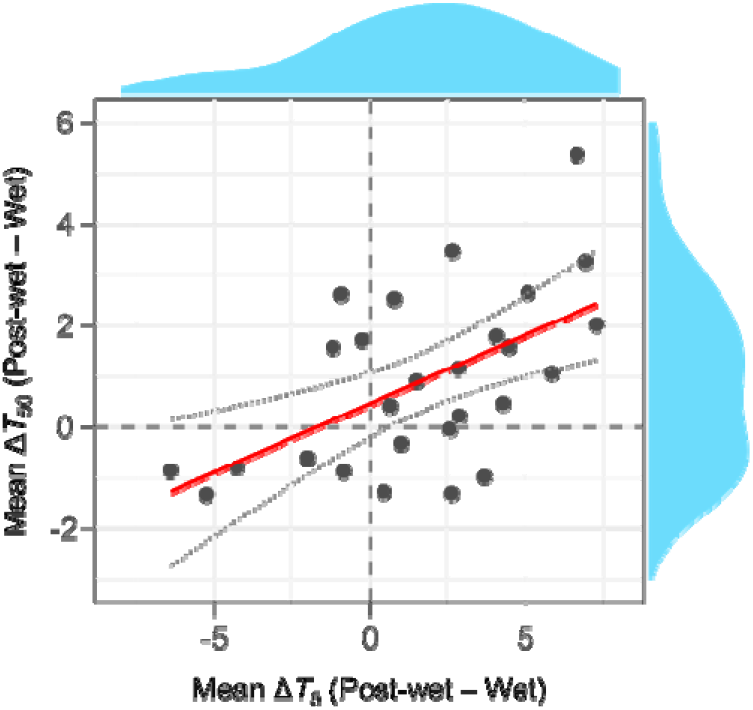
Coordinated seasonal shifts in *T*_5_ and *T*_50_ across 24 tree species in a seasonally dry tropical forest, Central Western Ghats. Species-mean Δ*T*_5_ (post-wet minus wet *T*_5_, °C) vs. Δ*T*_50_ (post-wet minus wet *T*_50_, °C) reveals positive correlation (*r* = 0.56, *p* = 0.0023), with most species showing parallel shifts (odds ratio = 7.3, *p* = 0.061). Standard major axis slope = 2.08, 95% CI: 1.49–2.90 (*R*^2^ = 0.32, *p* < 0.01).

### *T*_5_ ∝ decline width

SMA regression revealed a significant negative *T*_5_–DW relationship during both wet (slope = −0.61, 95% CI: −0.81 to −0.46, *R*^2^ = 0.50, *p* < 0.001) and post-wet (slope = −0.52, 95% CI: −0.70 to −0.38, *R*^2^ = 0.43, *p* < 0.001) periods, with identical slopes across seasons (χ^2^_1_ = 0.62, *p* = 0.43). Linear mixed-effects modelling confirmed the strong *DW* main effect (*F*_(1, 36)_ = 79.4, *p* < 0.001; marginal *R*^2^ = 0.59) but revealed habit-specific modulation during post-wet conditions (season × habit: *F*_(1, 28)_ = 4.71, *p* = 0.039; three-way: *F*_(1, 28)_ = 5.77, *p* = 0.023) (Figure 5). Thus, while high-*T*_5_ species consistently show narrow DW (prevention-focused), evergreens and deciduous species diverge in this trade-off under warming. This extends the prevention/forbearance framework to seasonal dynamics in a diverse SDTF community.

**Figure 5.**
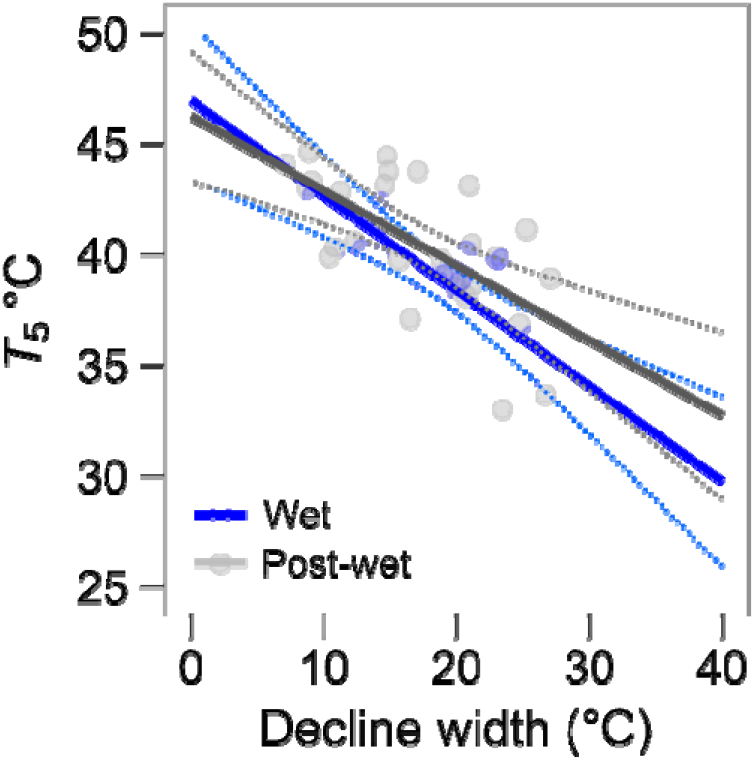
Relationship between temperature of PSII heat sensitivity initiation (*T*_5_) and decline width, the temperature window from *T*_95_ to *T*_5_. Standard major axis regression (linear) model fit was significant for both wet and post-wet periods the slopes were statistically indifferent.

### Proximity to PSII functional limits: thermal safety margin

Thermal safety margins based on *T*_50_ were large for all species, such that seasonal maximum air temperatures above the 90^th^ percentile never approached PSII damage midpoints in either period. In contrast, *T*_5_-based margins revealed that a few species operated closer to functional thresholds: during the wet period, *Saraca asoca* and *Schleichera oleosa* had *T*_5_ values near wet-season *T*_max_ > 90^th^ percentile, and during the post-wet period *Ficus macrocarpa* and *F. religiosa* showed similarly small margins relative to post-wet *T*_max_. Deciduous species generally maintained comfortable buffers, with none directly overlapping seasonal *T*_max_, although Careya arborea showed a reduced margin and trended towards the post-wet upper limits. Overall, most species retained substantial separation between *T*_5_ and realized air temperature extremes, but a small subset of late-successional evergreens and one early successional deciduous species already function near emerging seasonal heat constraints.

## Discussion

### Decoupling of phenological strategy and PSII thermal plasticity

A central assumption in tropical ecophysiology is that leaf phenological strategy and physiological thermotolerance regulation are coupled; i.e., evergreens investing in year-round PSII stabilisation, deciduous species relying on avoidance through leaf shedding (Sastry & Barua 2017; Sastry *et al*. 2018). Our results challenge this assumption directly. Both *T*_5_ and *T*_5_ increased modestly but significantly from the wet to post-wet period across the community, yet the magnitude of this acclimation was overwhelmingly species-idiosyncratic rather than structured by leaf habit or successional status. Physiological PSII regulation and phenological heat-avoidance thus appear to operate as largely independent axes of thermal strategy, deciduous species do not delegate heat tolerance to leaf shedding, and evergreens do not compensate for prolonged exposure through disproportionate PSII upregulation. That both groups acclimate to a similar degree suggests species draw on both strategies simultaneously, in combinations reflecting individual life-history trajectories rather than broad functional groupings.

Critically, our sampling captures this plasticity during a non-stress window, the wet and post-wet periods when leaves are fully mature, water is not limiting, and temperatures are rising but have not yet reached peak dry-season extremes. This design is deliberate and informative: by excluding the confounding effects of drought stress, leaf senescence, and defoliation that characterise peak dry-season measurements (Sastry & Barua 2017; Slot *et al*. 2021), we isolate the thermal signal in PSII plasticity. The modest but significant acclimation detected here, approximately 1.7°C in *T*_5_ and 0.9°C in *T*_50_ — therefore reflects genuine thermally-driven physiological adjustment rather than stress-induced or senescence-related changes in PSII integrity. This is consistent with the partial acclimation reported in urban-garden and field studies across diverse biomes (Tarvainen *et al*. 2022; Kullberg & Feeley 2024; Manzi *et al*. 2025), but extends those findings into a community-wide, paired, non-stress context that has not previously been quantified in SDTFs. The non-stress window thus provides a cleaner baseline for understanding the limits and mechanisms of PSII plasticity than peak-season comparisons, and we suggest it should be incorporated more systematically into future thermal acclimation studies.

### Deciduous species show higher *T*_50_ than evergreens: a leaf economics interpretation

The higher *T*_50_ observed in deciduous relative to evergreen species in our study contrasts with Sastry and Barua (2017), who reported the opposite pattern and interpreted it as reflecting greater constitutive membrane investment in evergreens exposed to chronic thermal stress. However, a critical contextual difference must be noted: their study was conducted in an urban garden setting where species were growing under modified microclimates, altered soil conditions, and reduced canopy competition, conditions that likely confound the expression of intrinsic thermotolerance strategies. Our study, by contrast, samples species in their natural forest context, where canopy position, interspecific competition, and seasonal microclimate interact to shape leaf traits as they would under evolutionary selection.

Within our natural forest community, we suggest the higher deciduous *T*_50_ reflects a leaf economics argument rooted in longevity and resource allocation. Deciduous species invest heavily in leaf construction during the brief wet-season window, producing leaves that must maximise photosynthetic returns within a constrained lifespan while tolerating rising post-wet temperatures before senescence. This compressed functional window may select for higher baseline PSII thermal robustness as an insurance strategy, not because deciduous species face greater chronic heat exposure, but because mid-season leaf replacement is not viable. This aligns with the slow-fast resource acquisition spectrum, where short-lived, high-return leaves in deciduous species are expected to show distinct thermal trait combinations relative to the more conservative investments of long-lived evergreen leaves (Russo & Kitajima 2016; Slot *et al*. 2021). Evergreen species, by contrast, can spread PSII damage and repair costs across longer leaf lifespans, tolerating moderate recurring thermal perturbation through D1 turnover and incremental acclimation rather than constitutive robustness (Yi *et al*. 2022). Direct comparison of leaf economic traits such as specific leaf area, leaf nitrogen, and leaf lifespan across both datasets would be a productive avenue for testing this interpretation.

### Conservation of the *T*_5_*–DW* prevention-forbearance trade-off across seasons

The negative *T*_5_ – *DW* relationship where species with higher thermal onset temperatures show narrower decline windows, was conserved across both seasons, confirming that the prevention-versus-forbearance axis represents a stable dimension of PSII thermal strategy rather than a seasonal artefact (Tiwari *et al*. 2021). Identical slopes in wet and post-wet periods indicate that as species shift their thermal onset upward with warming, the shape of their *F*_v_/*F*_m_ - temperature response is maintained. However, the habit-specific divergence in this relationship under post-wet conditions is noteworthy: evergreen and deciduous species separated along the prevention-forbearance axis as temperatures rose, suggesting that while the trade-off itself is conserved, the position species occupy along it is subtly modulated by habit under warming. This points to an interaction between phenological strategy and thermal response dynamics that warrants further investigation across the full annual cycle.

### Within-habit variation and the limits of binary phenological classification

Within both evergreen and deciduous groups, individual species showed considerable variation in the direction of seasonal change: some increased *T*_5_, *T*_50_, or *DW* from wet to post-wet, others showed no detectable shift, and a subset showed modest declines. This within-habit heterogeneity reinforces the conclusion that binary leaf habit classification captures only a coarse axis of thermal strategy variation, and that the species-specific nature of PSII plasticity documented here reflects genuine biological diversity that functional groupings cannot resolve. Fine-scale phenological attributes including the duration of mature leaf retention, the timing and pace of leaf senescence and emergence, and seasonal preferences in leaf turnover are likely to explain thermal strategies more precisely than a simple evergreen-deciduous dichotomy. Among the 27 species studied, meaningful differences in these phenological attributes exist that were not captured by our binary classification. For example, while most drought-deciduous species avoid the peak summer period by shedding leaves (E.g., *L. coromandelica, C. arborea, S. oleosa*) well before temperatures peak, a small number of species in this community retain or flush leaves during the later summer months (evergreens such as *A. heterophyllus, N. cadamba*, and *Ficus* spp.; deciduous species such as *T. bellirica, T. paniculata, T. anogeissiana*) (Nanda *et al*. 2012), which corresponds to one of the hottest periods of the year. Whether these species deploy distinct thermal tolerance strategies relative to their habit-group peers remains an open question that warrants dedicated investigation. Resolving these within-habit contrasts would require fine-frequency phenological monitoring across the full annual cycle, linking leaf stage and turnover rates directly to PSII thermal metrics measured at matched time points.

Beyond phenological complexity, resource allocation to leaf heat tolerance and structural leaf traits are expected to interact with the thermal strategies observed here. Investment in membrane thermostability, D1 protein turnover, and heat-shock protein synthesis all compete with investment in photosynthetic capacity and leaf structural robustness, implying that leaf economics should modulate both constitutive thermotolerance and its seasonal plasticity (Russo and Kitajima 2016; Sastry and Barua 2017). The species-driven variation in plasticity we observe is therefore likely to reflect, at least in part, differences in how species allocate resources across these competing demands, differences that are not captured by habit alone and that would require trait-level data to disentangle. While we did not collect leaf economic trait data in this campaign, this was a deliberate choice: our aim was to first establish whether phenological strategy and PSII thermal regulation are coupled or independent before introducing the additional complexity of morphological and chemical trait variation. Future studies integrating specific leaf area, leaf nitrogen, and leaf lifespan with the seasonal PSII framework developed here would provide a more complete mechanistic account of what drives species-specific plasticity in this community and would allow direct testing of whether the leaf economics argument proposed here to explain the higher deciduous *T*_50_ is supported by measured trait differences.

### Vulnerable species and implications for warming in SDTFs

Despite generally comfortable *T*_50_-based thermal safety margins across the community, consistent with findings from other tropical forests where *T*_50_ buffers remain large relative to ambient maxima (Slot *et al*. 2021; Kullberg & Feeley 2024); *T*_5_-based margins revealed a functionally significant vulnerable subset. Late-successional evergreens including *Saraca asoca* and *Ficus* species, alongside the early-successional deciduous *Careya arborea*, showed the smallest buffers relative to post-wet temperature maxima. That these species span both habit categories and successional positions reinforces the conclusion that vulnerability is not well-predicted by broad functional grouping. Their reduced margins likely reflect distinct mechanisms: *Saraca asoca* and *Ficus* species may reflect trade-offs of shade-adapted or strangler growth strategies (Schmidt & Tracey 2006) that prioritise carbon capture over thermal robustness, while *Careya arborea*’s fire-adapted open-canopy habit (Malasiya *et al*. 2022) may expose it to higher leaf temperatures than closed-canopy species. As seasonal maximum temperatures continue rising under climate warming, species currently operating near *T*_5_ thresholds face the greatest risk of crossing functional limits, not at peak dry-season extremes, but during the earlier post-wet warming transition that this study specifically captures. Trait-based frameworks that aggregate species into evergreen and deciduous functional types, risk masking this within-community heterogeneity, and our findings argue for species-resolved approaches to predicting thermal vulnerability in SDTFs.

## Conclusion

Across 27 co-occurring tree species in a seasonally dry tropical forest of the central Western Ghats, PSII thermal tolerance increased modestly but significantly from the wet to the post-wet period, tracking a approximately 4°C seasonal warming during a physiologically critical non-stress window. This plasticity was real but partial, consistent with the biochemical constraints on PSII acclimation documented across tropical systems, and insufficient to fully compensate for seasonal temperature shifts. Crucially, the magnitude of this acclimation was driven by species identity rather than leaf habit or successional status, with evergreen and deciduous species acclimating to a similar degree, directly falsifying the assumption that phenological strategy predicts physiological thermotolerance regulation. The prevention-versus-forbearance trade-off along the *T*_5_ and *DW* axis was stable across seasons, confirming it as a conserved dimension of PSII thermal strategy, while thermal safety margins based on *T*_5_ identified a functionally vulnerable subset of species including late-successional evergreens and one early successional deciduous species already operating near seasonal heat limits during the post-wet transition rather than at peak dry-season extremes.

These findings carry a direct message for trait-based approaches to predicting tropical forest responses to warming; leaf habit alone is an unreliable proxy for PSII thermal vulnerability, and models that partition species into evergreen and deciduous functional types will systematically misrepresent the heterogeneity of thermal strategies operating within a single community. Species-resolved assessments integrating leaf longevity, canopy position, and individual life-history context are needed to identify which trees are most exposed to emerging thermal constraints. As the post-wet warming transition intensifies under climate change, arriving earlier and reaching higher temperatures, the non-stress window characterised here will progressively narrow, and the species currently operating closest to *T*_5_ thresholds will be the first to experience functional heat limitation during what is currently their peak productive period.

## Supplementary Data

Supplementary Data D1: Sanskrit language version of the manuscript.

Supplementary Data D1: Kannada language version of the manuscript.

Supplementary Table S1: Species studied: Taxonomy, leaf habit, and ecological roles in Central Western Ghats seasonally dry Tropical forest

Supplementary Table S2: Summary of mixed⍰effects model results for thermal tolerance parameters.

Supplementary Figure S1: Post-wet and wet period plasticity in PSII thermal tolerance (*T*_5_ and *T*_50_) of co-occurring tree species across successional categories.

## Author contribution

RT conceived the research. RT and YK planned and designed the study. PTB, SNH, RGH, PN, JBM, SBS, MMM, SG, TN, AKS, VN, LBS, KTG, conducted thermal tolerance measurements and field data collection. YK and RT oversaw the field campaigns. NA, SBS, and MMM supported phenological characterization. RT collated and analysed the data and wrote the manuscript. All co-authors contributed to manuscript revision and approved the final version for submission.

## Acknowledgement

We thank Vasudeva HR for the help with the Sanskrit and Kannada language versions of the manuscript. RT and RM thanks the Wenner-Gren, Birgitta Sintring Foundations and the Swedish Vegetation Society for the postdoctoral funding at Uppsala. RT would like to thank Shāradā-Chandramaulishwara and Ubhaya Jagathgurus of Sringeri for their guidance.

## References

Bartoń K. (2018) MuMIn: multi-model inference. In: R package.

Bates D., Mächler M., Bolker B. & Walker S. (2015) Fitting linear mixed-effects models using lme4. Journal of Statistical Software 67.

Doughty C.E., Keany J.M., Wiebe B.C., Rey-Sanchez C., Carter K.R., Middleby K.B., … Fisher J.B. (2023) Tropical forests are approaching critical temperature thresholds. Nature 621, 105–111.

Dusenge M.E., Duarte A.G. & Way D.A. (2019) Plant carbon metabolism and climate change: elevated CO2 and temperature impacts on photosynthesis, photorespiration and respiration. The New Phytologist 221, 32–49.

Guan K., Pan M., Li H., Wolf A., Wu J., Medvigy D., … Lyapustin A.I. (2015) Photosynthetic seasonality of global tropical forests constrained by hydroclimate. Nature Geoscience 8, 284–289.

Harris G.C. & Heber U. (1993) Effects of anaerobiosis on chlorophyll fluorescence yield in spinach (Spinacia oleracea) leaf discs. Plant Physiology 101, 1169–1173.

Kattge J., Díaz S., Lavorel S., Prentice I.C., Leadley P., Bönisch G., … Wirth C. (2011) TRY – a global database of plant traits: TRY - A GLOBAL DATABASE OF PLANT TRAITS. Global Change Biology 17, 2905–2935.

Krause G.H., Winter K., Krause B., Jahns P., García M., Aranda J. & Virgo A. (2010) High-temperature tolerance of a tropical tree, Ficus insipida: methodological reassessment and climate change considerations. Functional Plant Biology 37, 890–900.

Krishnadas M., Kumar A. & Comita L.S. (2016) Environmental gradients structure tropical tree assemblages at the regional scale. Journal of Vegetation Science: Official Organ of the International Association for Vegetation Science 27, 1117–1128.

Krishnamurthy Y.L., Prakasha H., Nanda A., Krishnappa M., Dattaraja H. & Suresh H. (2010) Vegetation structure and floristic composition of a tropical dry deciduous forest in Bhadra Wildlife Sanctuary, Karnataka, India. Tropical Ecology 51, 235–246.

Kuhlgert S., Austic G., Zegarac R., Osei-Bonsu I., Hoh D., Chilvers M.I., … Kramer D.M. (2016) MultispeQ Beta: a tool for large-scale plant phenotyping connected to the open PhotosynQ network. Royal Society Open Science 3, 160592.

Kullberg A.T. & Feeley K.J. (2024) Seasonal acclimation of photosynthetic thermal tolerances in six woody tropical species along a thermal gradient. Functional Ecology 38, 2493–2505.

Malasiya D., Ray T., Verma S., Rajpoot R., Maurya D., Kumar T., … Khan M.L. (2022) Effect of forest fire on tree species diversity in the tropical dry deciduous forest of nauradehi wildlife sanctuary, madhya pradesh, central India. International Journal of Ecology and Environmental Sciences 48.

Manzi O.J.L., Mujawamariya M., Tarvainen L., Ziegler C., Andersson M.X., Dusenge M.E., … Uddling J. (2025) Photosynthetic heat tolerance partially acclimates to growth temperature in tropical Montane tree species. Plant, Cell & Environment 48, 7848–7861.

Nanda A., Prakasha H.M., Murthy Y.L. & Suresh H.S. (2012) Leafing phenology of canopy, under-storey trees and seasonality in a tropical evergreen forest of Bhadra Wildlife Sanctuary, karnataka, southern India. Indian Journal of Forestry 35, 457–462.

O’sullivan O.S., Heskel M.A., Reich P.B., Tjoelker M.G., Weerasinghe L.K., Penillard A., … Atkin O.K. (2017) Thermal limits of leaf metabolism across biomes. Global change biology 23, 209–223.

Pascal J.P. (1988) Wet evergreen forests of the Western Ghats of India. Institut francais de Pondichery, Pondichery.

Posch B.C., Amoanimaa-Dede H., Aparecido L.M.T., Atkin O.K., Bison N.N., Blonder B.W., … Hultine K.R. (2026) High-temperature acclimation of photosystem II in land plants. The New Phytologist 249, 1108–1123.

R Core Team, (2025) R: A language and environment for statistical computing.

Ramos D.M., Andrade J.M., Alberton B.C., Moura M.S.B., Domingues T.F., Neves N., … Cunha J. (2023) Multiscale phenology of seasonally dry tropical forests in an aridity gradient. Frontiers in Environmental Science 11, 1275844.

Russo S.E. & Kitajima K. (2016) The ecophysiology of leaf lifespan in tropical forests: Adaptive and plastic responses to environmental heterogeneity. In Tree Physiology. (eds G. Goldstein & L.S. Santiago), pp. 357–383. Springer International Publishing, Cham.

Sastry A. & Barua D. (2017) Leaf thermotolerance in tropical trees from a seasonally dry climate varies along the slow-fast resource acquisition spectrum. Scientific Reports 7, 11246.

Sastry A., Guha A. & Barua D. (2018) Leaf thermotolerance in dry tropical forest tree species: relationships with leaf traits and effects of drought. AoB Plants 10, x070.

Schmidt S. & Tracey D.P. (2006) Adaptations of strangler figs to life in the rainforest canopy. Functional Plant Biology 33, 465–475.

Schreiber U. (2007) Pulse-amplitude-modulation (PAM) fluorometry and saturation pulse method: An overview. In Chlorophyll a Fluorescence. pp. 279–319. Springer Netherlands, Dordrecht.

Schreiber U. & Berry J.A. (1977) Heat-induced changes of chlorophyll fluorescence in intact leaves correlated with damage of the photosynthetic apparatus. Planta 136, 233–238.

Silva J.O., Espírito-Santo M.M., Santos J.C. & Rodrigues P.M.S. (2020) Does leaf flushing in the dry season affect leaf traits and herbivory in a tropical dry forest? The Science of Nature 107, 51.

Slot M., Cala D., Aranda J., Virgo A., Michaletz S.T. & Winter K. (2021) Leaf heat tolerance of 147 tropical forest species varies with elevation and leaf functional traits, but not with phylogeny. Plant, Cell & Environment 44, 2414–2427.

Tarvainen L., Wittemann M., Mujawamariya M., Manishimwe A., Zibera E., Ntirugulirwa B., … Uddling J. (2022) Handling the heat - photosynthetic thermal stress in tropical trees. The New Phytologist 233, 236–250.

Tiwari R., Gloor E., da Cruz W.J.A., Schwantes Marimon B., Marimon-Junior B.H., Reis S.M., … Galbraith D. (2021) Photosynthetic quantum efficiency in south-eastern Amazonian trees may be already affected by climate change. Plant, Cell & Environment 44, 2428–2439.

Tiwari R., Hegde B., Bandaru P., Hegde S., Achar S., Muscarella R., … Gloor E. (2026) Tropical forest trees with higher thermal optima of photosynthesis exhibit lower PSII heat stability. Journal of Experimental Botany.

Tiwari R., Hegde B., Hegde S., Bandaru P., Ramesh Babu M., Somashekhara Achar K.G., … Gloor E. (2025) Contrasting seasonal variation of photosynthesis in evergreen and deciduous tree species from a tropical forest. Physiologia plantarum 177, e70410.

Wickham H. (2009) Ggplot2: Elegant graphics for data analysis, 1st edn. Springer, New York, NY.

Yamori W., Hikosaka K. & Way D.A. (2014) Temperature response of photosynthesis in C3, C4, and CAM plants: temperature acclimation and temperature adaptation. Photosynthesis Research 119, 101–117.

Yi X.-P., Yao H.-S., Fan D.-Y., Zhu X.-G., Losciale P., Zhang Y.-L., … Chow W.S. (2022) The energy cost of repairing photoinactivated photosystem II: an experimental determination in cotton leaf discs. The New Phytologist 235, 446–456.

